# Cell type-specific enrichment of somatic aneuploidy in the mammalian brain

**DOI:** 10.1101/2023.12.18.572285

**Authors:** Eran A. Mukamel, Hanqing Liu, M. Margarita Behrens, Joseph R. Ecker

**Affiliations:** Department of Cognitive Science, University of California, San Diego, La Jolla, CA 92037, USA; Genomic Analysis Laboratory, Salk Institute, La Jolla, CA 92037, USA; Computational Neurobiology Laboratory, Salk Institute, La Jolla, CA 92037, USA; Howard Hughes Medical Institute, Salk Institute, La Jolla, CA 92037, USA

## Abstract

Somatic mutations alter the genomes of a subset of an individual’s brain cells^1–3^, impacting gene regulation and contributing to disease processes^4,5^. Mosaic single nucleotide variants have been characterized with single-cell resolution in the brain^2,3^, but we have limited information about large-scale structural variation, including whole-chromosome duplication or loss^1,6,7^. We used a dataset of over 415,000 single-cell DNA methylation and chromatin conformation profiles across the adult mouse brain to identify aneuploid cells comprehensively. Whole-chromosome loss or duplication occurred in <1% of cells, with rates up to 1.8% in non-neuronal cell types, including oligodendrocyte precursors and pericytes. Among all aneuploidies, we observed a strong enrichment of trisomy on chromosome 16, which is syntenic with human chromosome 21 and constitutively trisomic in Down syndrome. Chromosome 16 trisomy occurred in multiple cell types and across brain regions, suggesting that nondisjunction is a recurrent feature of somatic variation in the brain.

---

Single-cell sequencing has revealed the diversity of brain cell types as defined by gene expression (scRNA-seq), chromatin organization (snATAC-seq) and DNA methylation (snmC-seq)^8^. While these assays focus on the diverse ways that cells express the information encoded in DNA, increasing evidence also points to diversity in the genetic sequence itself. Somatic mutations, including single nucleotide variants^2^, transposable elements^9,10^, and copy number variations (CNVs)^1,11^, have been detected in individual brain cells using both sequencing and *in situ* imaging. Recurrent variants can identify developmental lineages in healthy adults^12^, and rates of somatic variation have been linked with neurodevelopmental disorders, including autism^3,4,13^. However, because somatic variation is stochastic and present in a small subset of cells, the prevalence of mutations can be reliably assessed only by sampling large numbers of brain cells with sensitive whole-genome assays.

Single nucleus methylcytosine sequencing provides unbiased whole-genome coverage of sodium bisulfite-treated DNA^14,15^. By sorting single nuclei in individual wells and using liquid-handling robotics to efficiently scale library preparation, snmC-seq has been applied to samples of over 100,000 mouse brain cells^16,17^. The pattern of DNA methylation is a reliable marker of cell type identity that can identify fine-grained cell types aligning closely with transcriptomic and chromatin accessibility signatures^18^. Here, we took advantage of a large data resource generated using two technologies, single nucleus methylcytosine sequencing (snmC-seq2^19^) and multi-omic sequencing methylation and chromatin conformation (snm3C-seq^20^), which provide single base-resolution DNA methylomes with an average of 1.66 million reads per cell.

Because bisulfite sequencing samples the genome uniformly, we reasoned that the density of snmC-seq fragments across genomic regions could reveal variations in relative copy number^1,11^. Using snmC-seq data to detect CNVs and aneuploidies adds a new layer of information to existing data resources and directly connects structural variation with the fine-grained cell type identity defined by the DNA methylation pattern in the same cell.

## Single cell DNA methylomes identify chromosome duplications and deletions

We analyzed copy number variation using a large dataset of single-cell DNA methylomes from adult (P56) male C57BL/6 mice, covering most of the brain^17^. To estimate copy number in single cells, we analyzed the density of mapped DNA fragments in bins of size 100 kb or 1 Mbp, followed by normalization for local GC content and circular binary segmentation^21^. We focused on high-quality cells with at least 900,000 uniquely mapped reads and median absolute pairwise deviation (MAD) <0.3 (Fig. 1c). In total, the dataset includes 415,103 from 331 independent snmC-seq or snm3C-seq datasets^17^. We excluded genomic regions with low average coverage across all cells, including centromeres, repetitive sequences, and the Y chromosome (227 Mbp total).

**Fig. 1.**
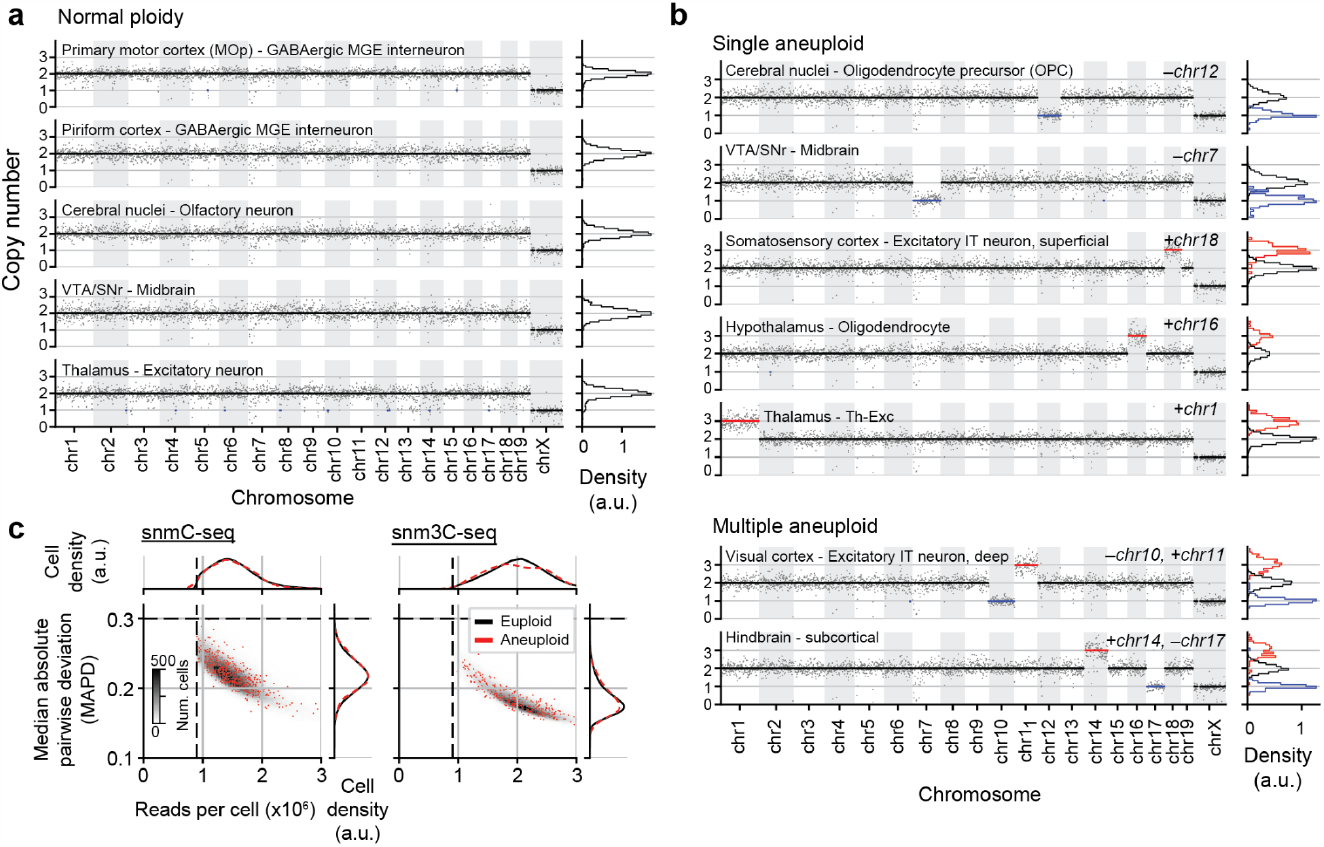
Aneuploid brain cells detected by single-cell DNA methylation sequencing. **a**, Examples of cells with normal copy number (euploid) from adult male mice assayed by snmC-Seq or snm3C-seq. Gray dots show normalized coverage in 1Mbp bins, and black lines show the estimated copy number. Cells are labeled with the region from which they were dissected and their cell class as determined by the DNA methylation^17^. Histograms (right) show normalized coverage distribution across bins; coverage on chrX is scaled by 2x. **b**, Examples of aneuploid cells with single or multiple duplications (red) or deletions (blue). **c**, Distribution of the number of reads per cell and the median absolute pairwise difference (MAPD) in normalized coverage between neighboring bins for cells from each assay. Dashed lines show the threshold for including cells. Aneuploid cells (red) have a similar distribution of both quality metrics compared with euploid cells (gray histogram).

The overwhelming majority of cells had a canonical pattern of coverage, with N=2 copies of each autosomal region and N=1 for chrX (Fig. 1a). A subset of cells had one or more whole chromosome loss or duplication events (Fig. 1b), consistent with previous observations^1^. We defined a cell as aneuploid if at least one chromosome was estimated to have a deletion or duplication over >90% of its extent (Extended Data Fig. 1c). By analyzing coverage with 100 kbp resolution, we found 734 aneuploid cells (0.175% of all cells, Fig. 1c). Increasing the bin size to 1 Mbp to reduce sampling variability (Extended Data Fig. 1a,b), we identified 1433 cells with a putative aneuploidy (0.349%).

Importantly, the quality of single-cell DNA sequencing data from aneuploid was equivalent to that of euploid cells, with similar distributions of total read depth and of median absolute pairwise deviation (MAPD) (Fig. 1c). The average number of sequenced reads was higher, and the MAPD lower, for cells assayed by snm3C-seq compared with snmC-seq. However, the percentage of aneuploid cells was the same for both modalities (Extended Data Fig. 1d).

### Recurrent somatic duplication of mouse chromosome 16

Losses and duplications were observed on all chromosomes (Fig. 2a). We identified more deletions of short compared with long chromosomes (Spearman r = –0.53, p=0.016), but duplications were not significantly correlated with chromosome length (p=0.11) (Extended Data Fig. 1e). In principle, some chromosome loss events could potentially be an experimental artifact caused by physical loss of a chromosome from the nucleus during the fluorescence activate nuclei sorting (FANS) procedure. Such artifacts can not explain chromosome duplication.

**Fig. 2.**
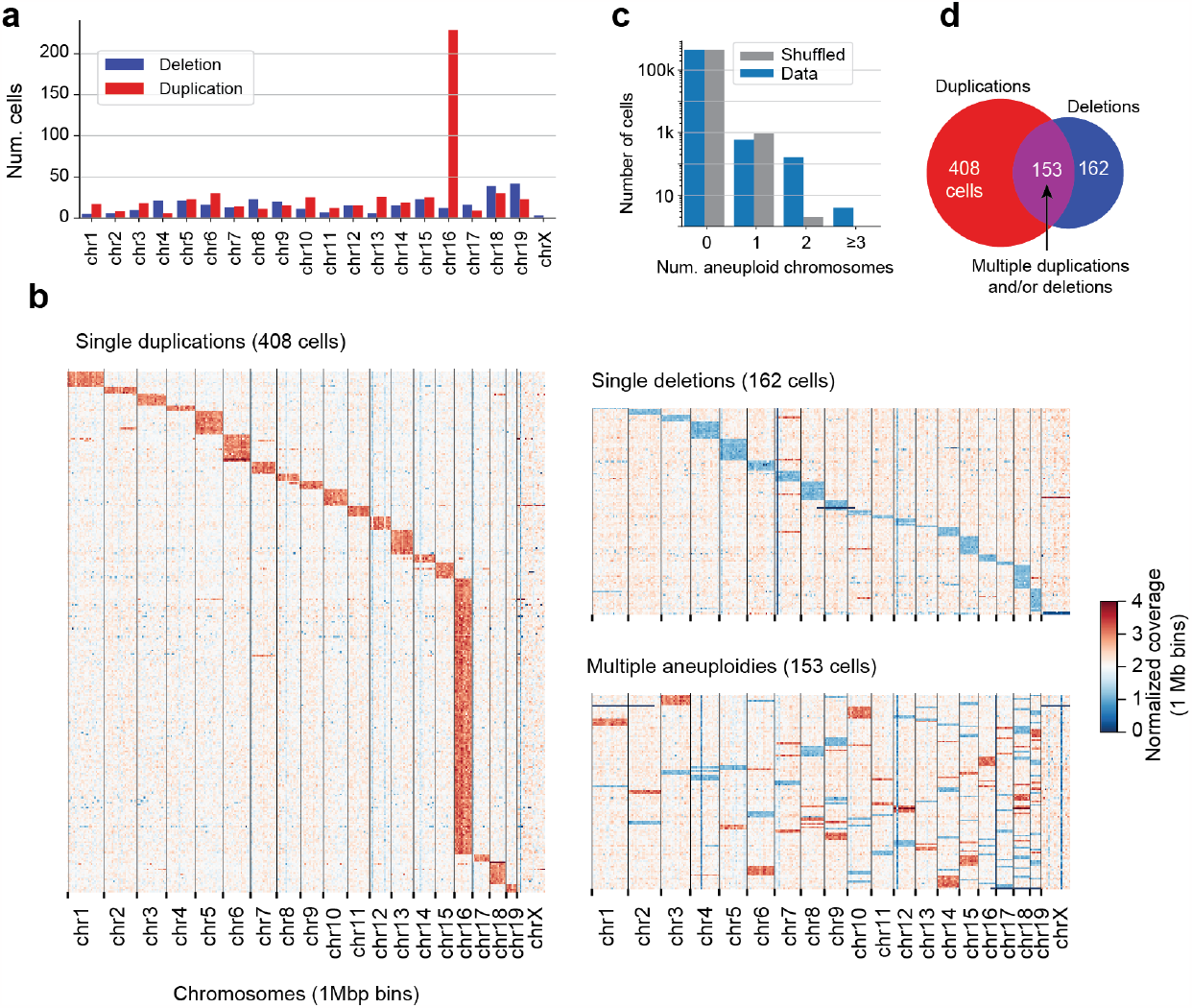
Enrichment of trisomy chr16 and multiply-aneuploid cells. **a**, Number of cells with a duplication or loss of each chromosome. **b**, Heatmaps showing the normalized coverage in 1Mbp bins for all aneuploid cells. Coverage in chrX is scaled by 2x. **c**, The number of cells with 2 or more aneuploid chromosomes was greater than expected for independent events (p<10^-300^, Binomial test). **d**, Almost 20% of aneuploid cells had both a deletion and a duplication.

We observed a striking enrichment of chromosome duplication (trisomy) on chromosome 16 (Fig. 2a,b). Trisomy 16 (T16) occurred in 234 cells, compared with an average of 18.3 cells with trisomy on other autosomes (13-fold enrichment, p<0.001, binomial test). The distal end of mouse chromosome 16 is syntenic with all of human chromosome 21^22^ (Extended Data Fig. 1f), and mice with trisomy of chr16 (e.g., Ts65Dn) have been used as a model of human Down syndrome (trisomy chr21)^23^. Notably, mouse chr16 and human chr21 contain the key oligodendrocyte lineage genes, *Olig1* and *Olig2*, and Ts65Dn mice have an increased proportion of GABAergic inhibitory neurons^24^.

Aneuploid cells were found in data from multiple independent cohorts of mice (Extended Data Fig. 1g,h). We identified at least one aneuploid cell in 229/331 mouse brain samples and ≥5 aneuploid cells in 40 independent samples.

Chromosome loss and duplication events were not independent across cells. We found 153 cells with 2 or more aneuploid chromosomes, 82-fold more than the expected number for independent events (p<10^-10^, binomial test, Fig. 2c,d). Most multi-aneuploid cells had 1 deletion and 1 duplication (131 cells), while 7 cells had ≥ 2 duplications and 15 cells had ≥2 deletions. The proportion of cells with multiple aneuploidies may be an underestimate, given our conservative criteria for calling aneuploidy. The pattern of correlated chromosome losses and duplications in individual cells is notable and suggests a potential mechanism involving the missegregation of chromosome pairs during mitosis.

### Cell type-specific atlas of brain somatic aneuploidy

Previous single-cell studies using whole genome sequencing identified somatic aneuploidy and CNVs in brain cells from mice^25^, humans^1,11,26,27^ and non-human primate^11^, but did not resolve the identity of the aneuploid cells. We took advantage of the DNA methylation information from snmC-seq data to assign cells to major classes and fine-grained cell types based on the CG and non-CG methylation pattern across the genome^17^. DNA methylation can distinguish cell types with high resolution, identifying 261 cross-modality annotated cell subclasses in our mouse brain dataset^17^. Because of the relatively small proportion of aneuploid cells, for this study, we used a coarse-grained cell type annotation with 48 major cell types (level 1 clusters), including 6 non-neuronal classes and 42 types of neurons (Fig. 3a). These cell types were distributed across 73 separately dissected annotated regions.

**Fig. 3.**
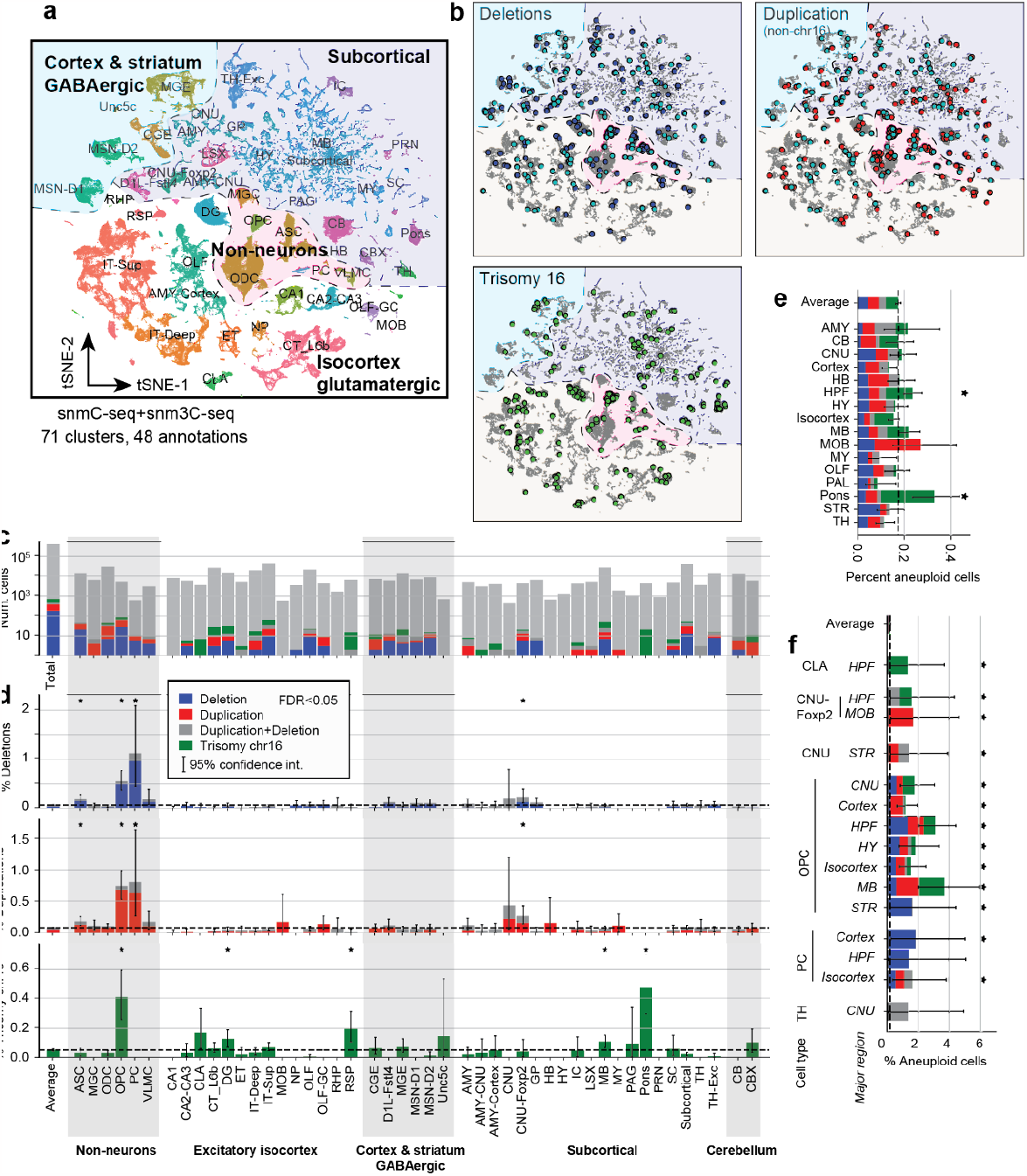
Aneuploid cells are enriched in oligodendrocyte precursors and pericytes. **a**, Embedding and annotation of all cells based on DNA methylation. **b**, Aneuploid cells are plotted on the embedding alongside euploid cells (gray), showing that aneuploidy is broadly distributed across cell types. Cyan cells have multiple aneuploidies. **c**, Number of cells of each major cell type, including euploid (gray) and aneuploid (colored) cells. **d**, Percent aneuploid cells of each cell type. Dashed line shows the average across all cell types. **e**, Percent aneuploid cells for each brain region. **f**, Percent aneuploid cells for each cell type and brain region that has at least 1% aneuploid cells.

Aneuploid cells were broadly distributed across neuronal and glial cell types (Fig. 3b-d, Table S2). Overall, 0.29% of neurons and 0.69% of non-neuronal cells had an aneuploidy, with slightly more chromosome losses than duplications. Among neurons, the rate of aneuploidy varied from 0.012% (CA1 pyramidal cells; 95% confidence interval=[0.0003–0.046%], beta distribution) to 0.47% (Pons, CI=[0.29-0.70%]). Glial cells had higher rates of aneuploidy, ranging from 0.11% [0.043-0.20%] for microglia (MGC) to 1.77% [0.89-2.94%] for pericytes (PC). Oligodendrocyte precursors (OPCs) also had a relatively high rate of aneuploidy (1.65% [1.33-2.00%]).

We used a binomial test to evaluate the enrichment of different types of aneuploidy in each cell type. Chromosome losses and duplications were significantly enriched (FDR<0.05) in three non-neuronal cell types (OPCs, PCs, and astrocytes, ASC), and in CNU-Foxp2 and Pons neurons.

Given the marked enrichment of T16, we tested whether this form of aneuploidy was specifically enriched in a particular cell type or brain region. Overall, 0.054% [0.046–0.060%] of neurons and 0.0088% of glial cells had T16. T16 was significantly enriched in OPCs (0.42%), as well as in the Pons (0.48%), midbrain (MB, 0.11%), dentate gyrus granule cells (DG, 0.13%), claustrum (CLA, 0.21%) and retrosplenial cortex (RSP, 0.16%).

## Discussion

Somatic mutations that alter the genome of a subset of an individual’s cells can impact brain function and contribute to disease processes, yet the extent of mosaic structural variation in mammalian brains has remained unclear. Studies based on *in situ* hybridization, spectral karyotyping techniques^7,28,29^, and more recently multiplexed FISH^30^, reported high rates of aneuploidy in human and mouse neurons ranging from ∼10 to ∼60% of cells. By contrast, single-cell whole genome sequencing (scWGS) yields far lower rates of aneuploidy in human neurons, from 0.5 to ∼5% of cells^1,11,25,26,31^. Although single-cell sequencing is considered a gold standard for measuring somatic mutations^32^, previous studies using scWGS have sampled only up to several hundred cells from a single brain region. Moreover, scWGS cannot identify the cell types of aneuploid cells.

Here, we increase the number of cells evaluated for aneuploidy by over ∼1,000 fold by taking advantage of single nucleus methylcytosine sequencing (snmC-seq) and multiomic (snm3C-seq) data from the BICCN covering the entire adult mouse brain^17^. We found that DNA methylome data can reliably and precisely identify aneuploidy in specific brain cell types.

Our data show that aneuploidy is rare in the adult mouse brain, affecting <1% of cells, in line with previous scWGS studies based on smaller numbers of cells^1,11,31^. However, we found an intriguing and non-random pattern of aneuploidy with recurrent trisomy of chromosome 16 and enrichment of aneuploidies in specific cell types, including Pons neurons and oligodendrocyte precursor cells.

Notably, mouse chromosome 16 is syntenic with human chromosome 21, which is constitutively trisomic in Down syndrome and has been reported to experience somatic duplications in the brain in the context of aging and neurodegeneration^33–36^. Mice with germline trisomy-16 die in utero^37^, but single brain cells with T16 have been reported previously^25^.

Single-cell DNA methylation sequencing data represent a novel opportunity for assessing somatic mosaicism in the context of epigenetically defined cell types. This approach can potentially be applied to DNA methylome and multi-omic sequencing from mouse^16,17^, human^38,39^, and other species^40^. The functional implications of aneuploidy could be assessed, for example, using multi-omic DNA methylation and transcriptome sequencing (snmCT-seq)^41^. Given the rarity of aneuploid cells in the brain, large-scale analyses of thousands of cells will be important to accurately define the landscape of somatic structural variation and determine its contribution to brain function and disease.

## Author contributions

EAM conceived the study and designed the research methodology. EAM and HL performed formal analysis. EAM prepared figures and wrote the original draft of the paper. EAM, HL, MMB and JRE reviewed the final draft.

## Data availability

Sequencing data used in this study are available from the NeMO archive (RRID: SCR_016152) with accession nemo:dat-m5wqy8t.

## Acknowledgments

We are grateful to Jingtian Zhou and Chongyuan Luo for their comments and to members of the CEMBA consortium who contributed to the DNA methylation dataset. This work was supported by grants from the Chan Zuckerberg Initiative Seed Networks for the Human Cell Atlas to EAM, and NIH/NIMH grant U19MH114831 to JRE. JRE is an Investigator of the Howard Hughes Medical Institute.

## METHODS

### Single cell DNA methylation data

All data used in this paper previously reported^16,18,42 17^. Experimental procedures using live animals were approved by the Salk Institute Animal Care and Use Committee under protocol number 18-00006. Samples were collected from adult male C57BL6 mice at postnatal day P56-P63. Nuclei were isolated using fluorescence activated nuclei sorting (FANS) to enrich NeuN+ neurons while retaining ∼10% non-neuronal cells. The library prepartion, sequencing, and mapping of data are detailed in the original publication^17^.

### Copy number estimation

Sequencing data were mapped using the YAP pipeline^16^. For each sample, we extracted the total coverage of non-CG cytosine positions within bins of size 100kb throughout the genome, excluding chrY. The coverage of cytosines is proportional to the number of sequencing reads mapped to each bin. We then used a modified version of ginkgo software^21^ to perform copy number estimation for each cell. First, we normalize the coverage throughout the genome by regressing on the GC content of each bin. Next, circular binary segmentation is used to split the genome into contiguous regions of uniform coverage density. The output of ginkgo analysis includes a table with normalized coverage for each cell in each genomic bin, and a second table with the estimated copy number for each cell in each bin. We also calculated the median absolute pairwise difference (MAPD) for each cell, defined as the median of the difference in normalized coverage between neighboring bins.

### Removing outlier cells

After visualizing the distribution of total reads and MAPD for each cell, we retained cells with ≥900,000 uniquely mapped reads (R1+R2) and MAPD<0.3. We further excluded outliers from each sample. We applied singular value decomposition (SVD) to transform the QC metrics (reads, MAPD) into uncorrelated principal components. We then excluded cells for which the absolute value of their principal component loadings was >3.

### Removing bad genomic bins

We removed genomic bins with low normalized coverage, corresponding to centromeric or repetitive regions. For each cell we calculated the normalized coverage (counts per million, CPM) in each bin. We averaged the CPM values across all cells in each sample, and multiplied the coverage on chrX by 2. We excluded bins with average CPM less than 24 or greater than 80.

### Statistical analyses

The rate of aneuploid cells (Fig. 3c-f) was estimated using a beta distribution. Given *k* aneuploid cells observed in a sample of *n* total cells, we determined the 95% confidence interval using the beta distribution, *Beta*(x; α = *k*, β = *n* − *k*) as implemented in scipy.stats.beta.interval^43^. Significant enrichment compared with the average rate of aneuploid cells was assessed using a binomial test. P-values from the binomial test were adjusted for multiple comparisons to control the false discovery rate (FDR) using the Benjamini-Hochberg (statsmodels.stats.multitest.fdrcorrection^44^).

### Analysis of DNA methylation in aneuploid cells

For each aneuploid cell, we identified a set of 10 euploid cells to serve as a control group for comparing the pattern of DNA methylation. Control cells were randomly sampled from the set of euploid cells that had the same major cell type label, sequencing technology (snmC-seq or snm3C-seq), and biological sample as the aneuploid cell. We calculated the difference of the aneuploid cell’s methylation level on the aneuploid chromosome from the average methylation (mCG or mCH) at each genomic bin for the group of control cells.

### Supplementary Table 1: List of aneuploid cells

**Extended Data Fig. S1.**
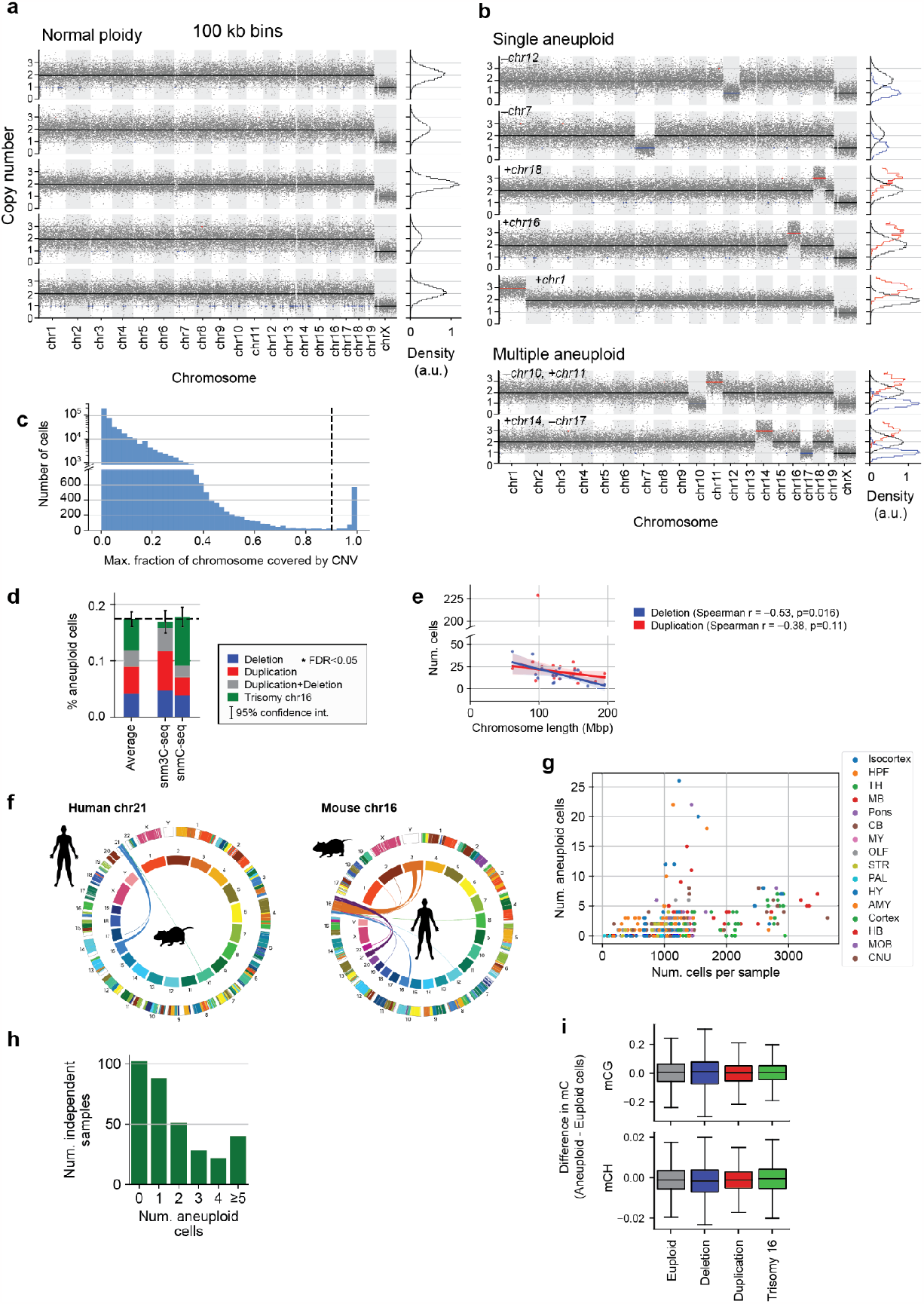
Aneuploid brain cells detected by single cell DNA methylation sequencing. **a,b** The same example cells shown in Fig. 1a,b are plotted here with a smaller bin size (100 kb). Gray dots show normalized coverage in 1Mbp bins, and black lines show estimated copy number. Cells are labeled with the region from which they were dissected and their cell class as determined by the DNA methylation^17^. Histograms (right) show the distribution of normalized coverage across bins; coverage on chrX is scaled by 2x. **c**, Histogram of the maximum fraction of a chromosome covered by a CNV for each cell. Dashed line shows the threshold (0.9) for defining a chromosome as aneuploid. **d**, Percent of aneuploid cells for each data modality (snmC-seq and snm3C-seq). **e**, Relationship between number of detected chromosome deletions or duplications and chromosome length. The lines show a robust regression fit with 95% confidence intervals. **f**, Synteny between human chromosome 21 and mouse chromosome 16, from the Jackson Labs synteny browser^22^. **g**, Number of aneuploid cells detected in each biological sample (derived from a pool of mice), vs. the total number of cells sampled. **h**, Number of aneuploid cells detected per sample. **i**, DNA methylation level on euploid vs. aneuploid chromosomes. Box plots show the difference in mCG or mCH between 100 kb bins on aneuploid chromosomes vs. the average of 10 euploid chromosomes in cells of the same type from the same sample.

